# Modelling Theta-Band Connectivity Between Occipital and Frontal Lobes: A Methodological MEG Study

**DOI:** 10.1101/123471

**Authors:** Darren Price, Matthew J. Brookes, Elizabeth B. Liddle, Peter F. Liddle, Lena Palaniyappan, Peter G. Morris

## Abstract

Measuring functional connectivity between cortical regions of the human brain has become an important area of research. Modern theory suggests that brain networks exhibit non-stationarity, constantly forming and reforming depending on task demands. A robust means of determining effective connectivity in the short-lived neural responses that occur in event related paradigms would allow the investigation of event related cortico-cortical dynamics. We present such a mathematical model of wave propagation, motivated by current neuroscience literature, and demonstrate the utility of the method in a clinical sample of schizophrenia patients. MEG data were acquired in 10 patients with schizophrenia and 12 healthy controls during a relevance modulation task. Data were filtered into the theta band (4-8Hz) and source localised using a beamformer. The model was implemented using Fourier analysis methods which uncovered an event related travelling wave moving from the visual to frontal cortices. The model was validated using Monte Carlo phase randomisation and compared with another plausible model of wave propagation in the cortex. Results from the clinical sample revealed that wave speed was modulated by task condition and patients were found to have less difference between conditions (ANOVA revealing a significant interaction between group and condition, p<0.05). In conclusion, our method provides a simple and robust means to investigate event related cortico-cortical brain dynamics in individuals and groups in the task positive state.

## Introduction

Measuring functional connectivity between spatially distinct brain regions has become an important goal for understanding human brain function. There is wide consensus that integration between functionally specific regions is crucial for the processing of information in the healthy brain (Buzsaki and Draguhn, 2004; Engel et al., 2001; Schnitzler and Gross, 2005; Uhlhaas and Singer, 2010). Moreover it has been hypothesised that a breakdown in the efficacy of such connections may underlie a number of neuro-pathologies including developmental disorders such as ADHD (Liddle et al., 2011), severe psychiatric illnesses such as schizophrenia (Friston, 1999; Phillips and Silverstein, 2003; Schnitzler and Gross, 2005; Stephane et al., 2008; Uhlhaas and Singer, 2010) and neurodegenerative disorders such as Parkinson’s disease (Hutchison et al., 2004; Schnitzler and Gross, 2005; Timmermann et al., 2004). Our ability to measure, with high spatio-temporal precision, the means by which functionally specific regions co-ordinate their activity is therefore of key importance. Brain network formation has primarily been investigated using functional magnetic resonance imaging (fMRI), and nearly two decades of research has demonstrated the existence of a relatively small number of robust large-scale distributed networks of connectivity; some associated with sensory systems (e.g. the visual and sensori-motor networks) and others associated with attention and cognition (e.g. the default mode and dorsal attention networks).

Using electrophysiological measurements, several authors have proposed that phase synchronisation of oscillatory activity serves as an effective means of cortico-cortical communication (Engel and Singer, 2001; Gray and Singer, 1989; Gray et al., 1989; Singer and Gray, 1995) in both short range (Eckhorn et al., 1988; Engel et al., 1991; Gray et al., 1989) and long range connections (Engel et al., 2001; Fries, 2005; Hipp et al., 2012; Salinas and Sejnowski, 2001; von Stein et al., 2000; Uhlhaas et al., 2010; Varela et al., 2001). Though useful, current methods primarily aim to identify aggregate brain networks observable over large time windows, in many cases lasting several minutes. However, human brain function relies on our ability to react to stimuli on a rapid timescale. This must necessitate the rapid formation and disintegration of transient brain networks, defined by short lived connections that are superimposed upon the distributed networks observable using traditional static metrics (Engel et al., 2013). fMRI is an indirect measure of brain activity based upon slow haemodynamics and thus is not well suited to the investigation of these transient brain states.

We aimed to use the relatively short-lived induced envelope from a seed location, and predict activity at other brain regions using a transfer function. This transfer function takes into consideration the non-linear effects of temporal delay, thereby capturing the transient effective connectivity associated with a particular cognitive task. The model presented here is informed by relevant knowledge of neuronal functioning showing that oscillatory coupling between regions results in effective connectivity and cortical communication.

Communication between cortical regions may be achieved by synchronised firing of neuronal populations, since the integration of such coincident firing at the target population increases the likelihood of a neuronal response (Abeles, 1982; König et al., 1996; Salinas and Sejnowski, 2001). Therefore, tightly synchronised oscillations are more effective than non-synchronised outputs at transferring information from one region of the cortex to another (Alonso et al., 1996; Azouz and Gray, 2000; Bruno and Sakmann, 2006). Engel et al. (2013) have described short lived spatiotemporal patterns of connectivity as intrinsic coupling modes (ICMs) categorised loosely by two mechanisms of interaction; the first mechanism involves a linear fixed phase relationship between the oscillations generated by spatially distinct brain areas (Gow Jr et al., 2008; Gross et al., 2001; Ioannides et al., 2000; Jerbi et al., 2007; Nolte et al., 2004; Schlögl and Supp, 2006; Schoffelen and Gross, 2009; Tass et al., 1998). The second involves temporal correlation between the amplitude envelopes of neural oscillations in different regions (Brookes et al., 2011a, 2012; Bruns et al., 2000; Hall et al., 2013; Hipp et al., 2012; Liu et al., 2010; de Pasquale et al., 2010). Both forms of coupling identify networks of functional connectivity in the human brain. Furthermore, both forms have a degree of spatial concordance with the networks observed using fMRI (Brookes et al., 2011a; Marzetti et al., 2013; de Pasquale et al., 2010; Schölvinck et al., 2010).

However, the precise mechanism by which amplitude correlation occurs is not known. One mechanistic interpretation is that state changes in distributed cortical networks are modulated on slow timescales by neuro-modulatory input (Leopold et al., 2003; Munk et al., 1996). However, on shorter timescales, i.e. during a cognitive task containing short trials, amplitude modulation is likely to be driven by transient synchronisation of local cortical oscillations (Neuper et al., 2006; Pfurtscheller and Aranibar, 1977; Pfurtscheller and Lopes da Silva, 1999). The true picture is likely a combination of the two. However, for building our model, we will assume that transient connections between cortical regions during a task are driven by inter-cortical communication. When two or more neuronal populations are engaged in coherent activity, phase differences between sending and receiving populations may regulate the flow of information (the effective connectivity) (Fries, 2005; Womelsdorf et al., 2007) a proposition that forms the basis of effective connectivity measures in the EEG and MEG such as dynamic causal modelling (DCM) (Friston et al., 2003; Stephan et al., 2007). In the stimulus induced response this directed form of connectivity may also manifest in the envelope as correlation of event related synchronisation (ERS), even in the absence of long range phase coherence (Bruns et al., 2000). Therefore, inter-cortical delays in transient ERS and event related desynchronisation (ERD) might provide a phase independent marker for effective connectivity. Importantly, this theory implies that greater synchronisation corresponds to communication that is more efficient. Locally, this should be evident in the envelope as more rapid ERD and ERS in the post stimulus response (in other words a more rapidly modulated envelope in both communicating regions). In terms of cortico-cortical communication, this increased efficiency should result in a decrease in delay. Therefore, we make the prediction that temporal delay between cortical sites is proportional to the modulation frequency of the envelope.

In order to test a model based on these assumptions it is necessary to adopt a paradigm known to elicit long-range communication, and to elicit reliable components in the stimulus-induced response. The theta band has been implicated in communication between the occipital and medial frontal regions during working memory (Sarnthein et al., 1998) and visual attention paradigms (Gregoriou et al., 2009), suggesting a role for theta in long range communication between distant cortical sites. Furthermore, the amplitude of post stimulus theta ERS scales with task difficulty (Brookes et al., 2011a, 2011b; Jensen and Tesche, 2002) supporting the idea that theta ERS is somehow functionally significant in visual working memory. Anatomically, the visual processing pathway is comprised of a cortical hierarchy that sub-serves cognitive function by participating in both feed-forward and feedback cortical connections. Feed-forward signals convey sensory information, while feedback mechanisms provide modulatory input (Bastos et al., 2015a, 2015b; Felleman and Van Essen, 1991; Markov et al., 2014). Furthermore, sensory-guided goal-directed behaviour necessary for the completion of most cognitive tasks, relies on both feed-forward (Brookes et al., 2011a; Donner et al., 2009; Glimcher, 2003; Mazaheri et al., 2009, 2010) and feedback networks (Donner et al., 2008, 2009; Driver et al., 2009; Gold and Shadlen, 2001; Horwitz et al., 2004; Kastner and Ungerleider, 2000; Kim and Shadlen, 1999; Moore and Armstrong, 2003; Nienborg and Cumming, 2009; Ress and Heeger, 2003; Serences and Yantis, 2006; Zanto et al., 2011). Bastos et al. (Bastos et al., 2015a) have recently shown that, in the primate visual system, feed-forward and feedback influences are carried by distinct frequency ranges of neural oscillations. Specifically, theta (∼4Hz) and high gamma band (∼60-80Hz) oscillations were involved in routing of information from V1 to higher processing areas, while feedback influences were carried by beta (∼14-18Hz) oscillations. Inter-regional delay of power fluctuations between several functional networks have also been observed during a visual working memory study in humans (Brookes et al., 2011a; Liddle et al., 2016) suggesting that the relevance modulation task is a useful paradigm for investigating striatal-frontal connectivity. It was found, in both of these studies, that trial averaged induced responses revealed a burst of theta oscillations peaking at ∼200ms post stimulus onset in the visual cortex. A similar theta waveform was observed in the frontal regions, but was delayed relative to the stimulus by ∼400ms. If this observable theta burst is indeed representing a feed-forward influence within a hierarchical network it follows that a direct correlation should be observable between the theta waveform within these separate regions, if the mapping function between those two regions can be determined accurately.

Here, we aim to investigate the precise relationship between visual and frontal theta-band envelopes. Using MEG, we measure the theta band waveforms that are induced across multiple brain locations by a cognitive stimulus. We formulate two simple models of propagation of the theta envelope waveform from the occipital cortices to anterior brain regions. In the first model, we assume that wave components at all frequencies travel through the brain with the same velocity, such that the theta waveforms measured in the frontal regions are simply time-delayed reconstructions of the equivalent theta waveform in the visual cortex. In the second model, we employ a model to test the prediction that higher frequency envelope modulations will travel through the brain with a shorter delay. As a result, the envelope modulation frequency will be linearly related to the propagation delay, leading to a temporally distorted theta waveform in distant cortical regions. Note that this is not the same as measuring phase lag in the carrier signal. We present evidence that the latter model offers a useful means to investigate connectivity between distal brain regions (specifically visual to frontal). We also show that the parameters characterising the model are affected by the cognitive task undertaken by the subject. Finally, we show that these same parameters differ in patients with schizophrenia compared to healthy controls.

## Theory

In order to assess connectivity between two distal brain areas we first begin with a simple description of the two models to be tested. Consider first a simple sinusoidal waveform, induced by an external stimulus (e.g. visual stimulation) within a neural network at brain location 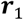. This waveform is subsequently “transmitted” via some means to a spatially separate brain location, 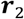 (see Figure 1A for an example).

**Figure 1:**
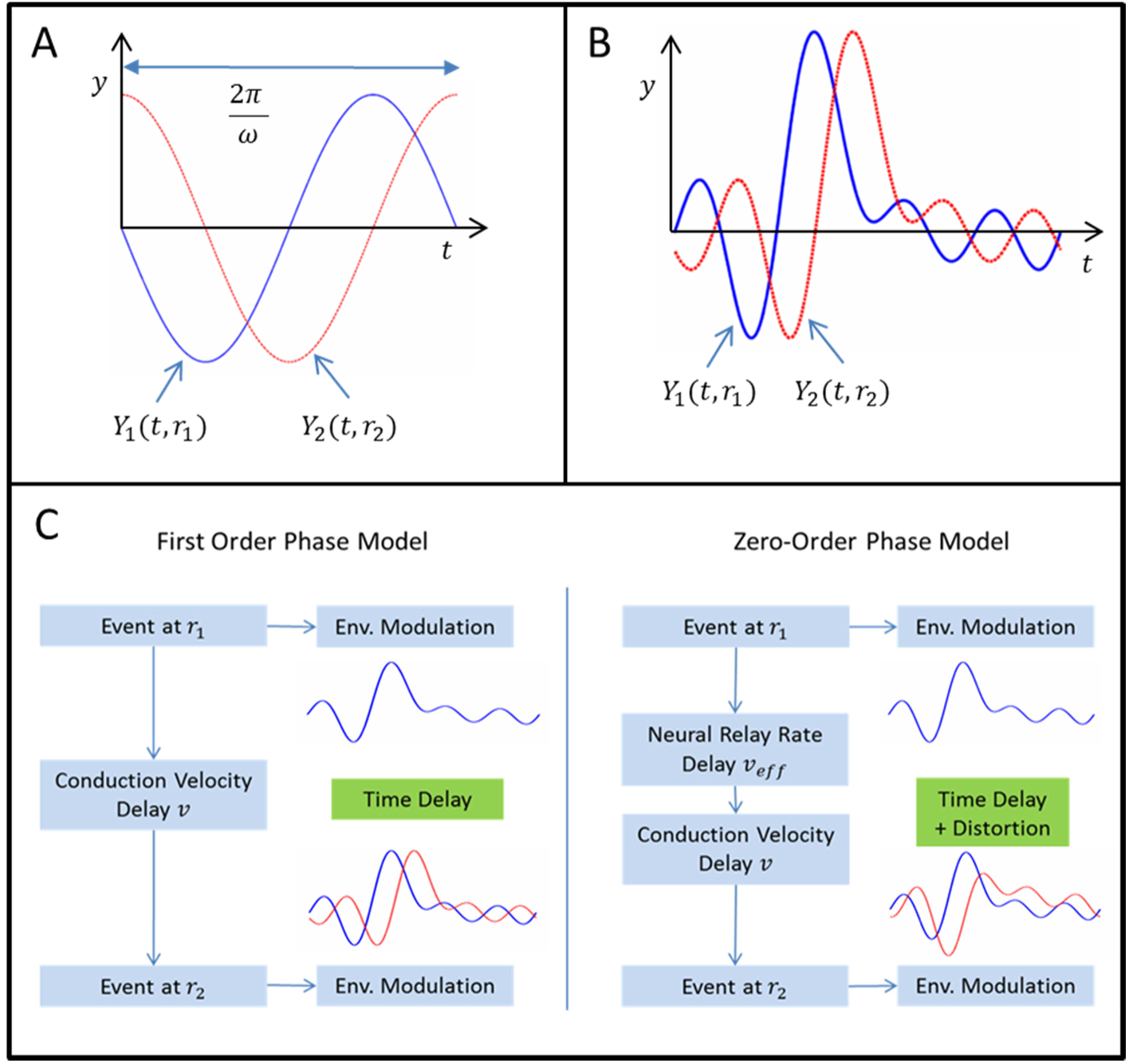
Two simple models of conduction through the brain. A) A phase shifted sinusoid. B) First order phase shift representation of a waveform with multiple frequency components, leading to a time shifted version of the original data. Note that the end of the original data is appended to the beginning of the time shifted data. D) Flow charts showing the difference between the zero-order and first order models.

If 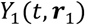 represents the measured waveform at location 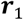 then taking the simplest possible waveform, we can assume

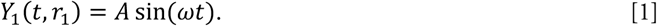

Note that amplitude, *A*, and frequency, *ω*, are determined by the external stimulus driving the waveform (i.e. if the stimulus were a flashing light, then *ω* would likely be defined by the frequency of the flash) and the localised cortical response to that stimulus at brain location 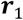. Now assume that 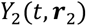 is the transmitted waveform measured at brain location 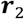. Note that the signal has, via some unknown path, been transmitted between 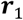 and 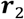 over a distance *δx*, with some effective velocity *υ*. Mathematically we guess that 
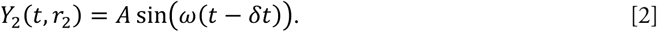

Here, *δt* is the time taken for the wave to travel from 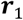 to 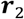 and so

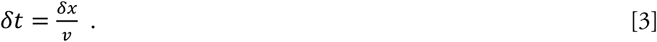
Therefore,

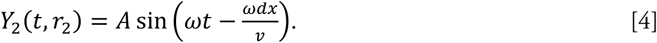

Note here that unlike the general case for a physical wave (e.g. light waves), the parameters *ω* and *υ* are unrelated. *ω* is likely to depend on the stimulus and response function of brain area 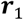 whereas *υ* represents conduction velocity in the brain.

The above treatment only accounts for a pure wave of a single frequency component. To generalise this, consider what happens if multiple different frequency components originate at 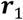. Here,

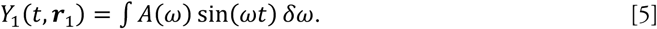

Again, assuming that a linear shift in the waveform between *r_1_* and *r_2_* we could write that

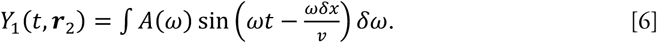

Note that *A*, the amplitude of each frequency component is allowed to vary with frequency. Recall also that for our simple case, *δx* and *υ* are constant. We now rewrite the expression for *y_2_* as

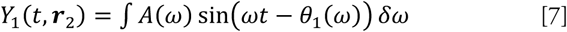
Where

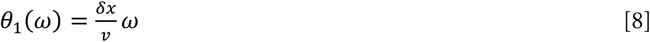

In words we state that *θ*_1_ is a phase term which is directly proportional to frequency *ω*. Implicit in the model is that we assume that conduction velocity, *υ*, is frequency independent, meaning that the measured waveform at 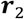 is simply a time shifted version of that measured at 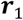. This simple model is depicted in Figure 1B and, since *θ*_1_ is a first order polynomial in *ω*, we term this a *first order phase* shift model.

The assumption inherent in the first order phase shift model is that all amplitude modulation frequencies travel with the same velocity, *υ*. However, alternative models may be proposed where other factors affect propagation velocity. One possible factor is the synchrony of the cortical population. Here we assume that a rapid modulation of band-limited power reflects a rapid change in synchrony of the local neural population. Neurons within the local population entrain distant connected neurons resulting in a rapid change in power at the connected region. Therefore both regions will show rapid modulation of power with a time delay that is proportional to the modulation frequency. This effect is independent of axonal conduction velocity which is assumed here to be relatively fast and of constant velocity. On different trials, and within different neuronal subpopulations, amplitude modulations of varying frequency add linearly resulting in a more complex average waveform.

For a simple idealised envelope with sinusoidal time course:

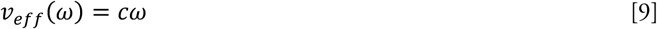
Where c is an unknown constant. Note that we do not suggest that physical conduction velocity through white matter changes with wave frequency, rather we use *υ* as an effective velocity which accounts for relative delay in action potential generation caused by different group delay. To denote this, we employ the term *υ_eff_*. Substituting Equation [9] into the above model we can write:

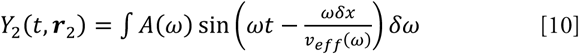
Or simply:

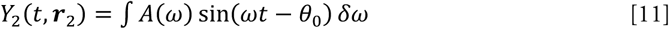
Where *θ_0_* = *δx/c* and is a constant for all frequency components. In other words, the phase shift is independent of frequency (i.e. *θ*_0_ is a zero-order polynomial). We term this model a zero-order phase shift. In fact, the zero order phase shift results not in a time shifting of the original waveform as depicted in Figure 1B, but rather a distorted version of the original waveform. This depicted in Figure 1D which shows two flow charts detailing the first order and zero-order phase shift models of transmission between two brain regions.

## Materials and Methods

### Participants and Paradigm

MEG data were acquired in 22 subjects. This comprised 10 patients satisfying the DSM-IV criteria for schizophrenia (Leckman et al., 1982)(Liddle et al., 2002), and 12 healthy controls not differing significantly from the patient group in mean age, sex or parental socioeconomic status (Pevalin and Rose, 2002). Nine patients (75%) were receiving anti-psychotic medication. Patients below the age of 18 or over 50, or with an IQ score of below 70 on the Quick Test (Ammons and Ammons, 1962), were not considered for the study. Healthy controls were recruited from the community via advertisements and were free from Axis I psychiatric disorders. The study was given ethical approval by the National Research Ethics Committee, Derbyshire, United Kingdom. All participants gave written informed consent.

All subjects took part in a “Relevance Modulation” task designed specifically to modulate the relevance to the subject of incoming visual stimuli. Note that this task has been employed in previously published studies (Brookes et al., 2011a) and we include its description here again for completeness. The subject was shown a series of visual stimuli (duration=800ms) in which pictures of butterflies were alternated with pictures of ladybirds. Inter stimulus intervals (ISI’s) were jittered, a single stimulus being presented, on average, every 2s. A single block comprised presentation of 40 alternating stimuli, followed by a resting phase. Blocks were 120s in total. At the beginning of each block, subjects were informed whether the *relevant* stimuli were the butterflies or the ladybirds. In blocks where butterflies were relevant, the subject’s task was to respond when the butterfly on the screen matched an example butterfly (shown at the start of the block) on three dimensions (shape, outer wing colour; inner wing colour); in blocks in which the ladybirds were relevant (LB blocks), the subject’s task was to respond if the number of red ladybirds in the picture was equal to the number of yellow ladybirds. The number of ladybirds ranged from 4 to 6, and they were presented in pseudo-random positions and orientations. In both block types, the “target” stimuli were rare (probability = 0.05), and these “target” trials were excluded from analyses so as to avoid confound from a motor response. Eight blocks were presented to each subject, in the order: BF, LB, LB, BF, LB, BF, BF, LB. Both task types (responding to a butterfly that matched the example; responding to equal numbers of red and yellow ladybirds) were designed to require close attention to the relevant stimulus.

### Data Acquisition and Preprocessing

MEG data were acquired using the third order synthetic gradiometer configuration of a 275 channel MEG system (MISL, Coquitlam, Canada), at a sampling rate of 600Hz and using a 150Hz low pass anti-aliasing filter. Three head position indicator (HPI) coils were attached to the subject’s head at the nasion, left preauricular and right preauricular points. These coils were energised with a subject inside the scanner to allow localisation of the head relative to the geometry of the MEG sensor array. The surface shape of the subject’s head was then digitised using a 3D digitiser (Polhemus, Isotrack) relative to the coils. The surface shape of each subject’s head and the coil locations were then coregistered with an anatomical MR image, acquired using a Philips 3T Achieva MR system at 1×1×1mm^3^ isotropic resolution using an MPRAGE sequence.

In order to project sensor space MEG data into brain space a scalar beamformer spatial filtering approach was employed (Robinson and Vrba, 1999; Van Veen et al., 1997). Beamformer weighting parameters were computed on a regular 8mm grid spanning the whole brain space. A different covariance matrix was computed for each frequency band. The forward model was based upon a multiple local sphere head mode (Huang et al., 1999) and the dipole model described by (Sarvas, 1987). Multiplication of the theta band filtered MEG data by the beamformer weights generated a timecourse of electrical activity for each voxel. Theta-band envelope time-series were then generated Hilbert transformation. These envelopes were averaged across all of the relevant and irrelevant trials in the dataset in a time window spanning *0s* < *t* < *2s* where *t* represents peri-stimulus time.

### FOPS and ZOPS Model Generation

Our primary aim, as detailed in our introduction, was to test whether the zero order phase shift (ZOPS) model was better able to measure transmission of a waveform through the brain than the first order phase shift (FOPS) model. To do this, we first take the measured waveform, 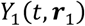 at brain region 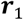 and act upon it using simple Fourier analysis to simulate the waveform at 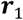. The simulated waveform can then be compared to the measured waveform in order to test which of the two models best fits the data. We denote the Fourier transform of 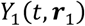 as 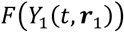) which can be written:

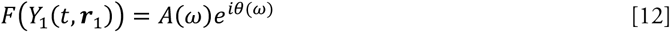
Where, *A*(*ω*) represents the amplitude and *θ*(*ω*) the phase of each frequency component in Fourier space. We now simulate the signal at 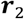 based upon the signal from *r_1_*, using the FOPS model as:

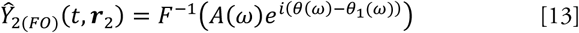

Note here *F^-1^* represents the inverse Fourier transform, and we use the ‘hat’ notation 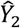 to represent the simulated version of *Y_2_*. Note also that as required for the FOPS model, *θ*_1_(*ω*) = *pω* where p is a constant. Similarly it is possible to simulate the signal at 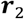 based upon the signal from *r*_1_, using the ZOPS model as:

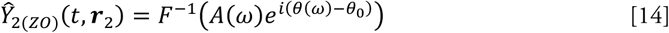
Where *θ*_0_ is a constant, meaning that to generate the simulated ZOPS signal we modify the phases of each frequency component by the same amount, irrespective of frequency.

#### Model testing and statistical analyses

In order to compute the similarity between the model and the data, correlation coefficients were computed between the measured theta timecourse *Y_2_* and the modelled theta timecourse 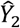. In order to generate the model, a time series, *Y*_1_, was taken from a seed location in primary visual cortex and used, alongside either Equation 13 or 14, to generate the modelled timecourses 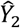.

For the zero order phase shift, *θ*_0_ was allowed to vary, taking 100 equally spaced values in the range [0 < *θ* < 2π]. A set of 100 correlation coefficients was calculated for every voxel in the brain and in all cases the highest correlation across the 100 phase values was extracted and used in further analysis (i.e. we select the phase shift value that best fits the data for any one single voxel).

For the first order phase shift, the phase change effectively amounts to a shift in time of the original seed time series. 100 iterations of the model time series were therefore computed with the time shift equally spaced between 0s and 2s. Again a set of 100 correlation coefficients was calculated, for every voxel in the brain, and the highest correlation across the 100 time shift values was extracted (i.e. in this case we select the time shift value that best fits the data for any one single voxel).

A potential limitation of the correlation approach is that for MEG, signals derived from multiple spatially separate voxels are not necessarily independent. This “signal leakage” between voxels is a direct result of the ill posed MEG inverse problem and is particularly problematic for voxels close to the centre of the head. In such cases the signal to noise ratio is very low and the projected timecourse can often represent a heterogeneous mix of the cortical signals. For this reason, in addition to correlation coefficients we also computed covariance. Computing the covariance between the model and the data not only accounts for similarity between *Y*_2_ and 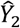 but also the variance in *Y*_2_. In beamforming, the variance of the signal is downweighted according to the norm of the weights vector (see Hall et al 2013) meaning that in deep regions, where leakage is high, the variance of 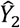 will be relatively low. We would therefore hypothesise that the covariance metric would generate a more accurate spatial map than the correlation metric.

To generate the correlation and covariance images, each subjects individual MRI was segmented and the brain extracted using the brain extraction tool (BET2) in FSL (Smith, 2002). The brain extracted MRI for each subject was then coregistered to MNI space using FLIRT (Jenkinson and Smith, 2001) and the same transform was applied to the functional (beamformer derived) data which were averaged across subjects in standard space. The covariance and correlation maps were then computed on an 8mm grid in MNI space using the subject averaged data. All covariance and correlation maps were interpolated to a 1mm^3^ isotropic MNI grid, overlaid on to a standard MNI brain (MNI152 1mm brain, available in FSL) yielding 100 covariance and 100 correlation maps (i.e. one map was available for each of the 100 phase shifts (in the case of ZOPS) or time shifts (in the case of FOPS). Animations were generated by sequencing these images from low to high phase shift. Finally, maps of optimum phase for every voxel in the brain were generated.

Calculation of statistical confidence intervals for each method, and comparisons between methods, were approximated by Monte Carlo phase randomisation of real signals extracted from the control subjects. Firstly, timecourses were extracted using beamformer source localisation at 50 locations spanning the visual and frontal cortices, chosen to be representative of the whole head. For each iteration of the Monte Carlo simulation, the extracted signals were phase randomised in the Fourier domain using the same method as described in (Prichard and Theiler, 1994) to maintain the covariance structure between locations, and preserving zero phase lag correlations such as those caused by weights correlation (see Fig 1.02). Each surrogate dataset was analysed using the same methodology as that used in real data. The phase-randomisation process was repeated 1000 times for each location producing a null distribution of *r* values. All values from >90mm were grouped to produce a general null distribution.

**Figure 1.02.**
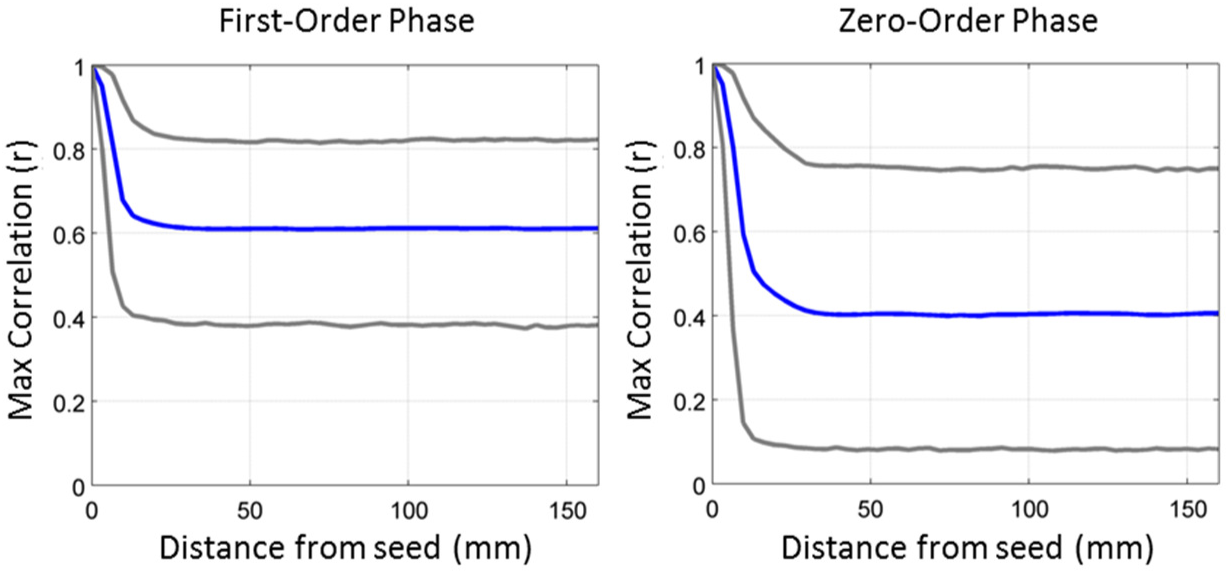
Results generated by phase randomisation simulations. The mean correlation of the null distribution is shown in blue, and the lower and upper 95% confidence limits are shown in grey below and above the mean. The effect of seed blur on the envelope correlation can be observed near 0mm. However, this flattens off at >30mm after which point the mean remains relatively constant. Note that FO method generates consistently higher mean correlation values under the null distribution than the ZO method. This difference needs to be taken into account when comparing procedures in real data.

### Between Groups Comparisons (Clinical Data)

For the between groups comparisons of the controls and clinical sample, analyses were performed on individual subjects. Here the delay estimates for each model were calculated per voxel, and then averaged across distance bins producing 9 delay values (1 per distance bin) for each subject. These values were then checked for statistical normality. Our hypothesis was focused on the frontal cortex, so only the last two distance bins (encompassing the frontal cortices >120mm from the seed) were entered into an ANOVA model (2 × condition, 2 × group, 2 × distance).

## Results

### Healthy Volunteer ZOPS Connectivity Analysis

Figure 2 shows results of our ZOPS connectivity analysis applied to data from healthy subjects, with a seed location in primary visual cortex. Analysis was limited to the averaged theta (4-8Hz) envelope timecourse. Figures 2A and B show axial and sagittal correlation images for phase values varying from 0° to 91°; Figure 2A shows the case for the relevant stimuli whilst Figure 2B shows the case for irrelevant stimuli. (The locations of the axial and sagittal slices used are shown in Figure 2H.) It is noteworthy that the ZOPS model generates high correlation, with the voxels exhibiting the highest correlation appearing to move anterior, as phase value is increased. The peak correlation of r=0.97 was located in medial frontal cortex and p values (obtained using the Monte-Carlo phase randomisation method) survive a Bonferroni correction for all 4626 voxels tested (p_correct_ed<0.05). This was considered a very conservative significance test. The anterior movement of regions of high correlation between the data and the ZOPS model, as phase is incremented, generates the effect of a ‘travelling wave’ propagating through the head. Note that the correlation values are affected by task condition, with high correlation for relevant stimuli and lower correlation for irrelevant stimuli. Figure 2C and D shows the equivalent covariance images, derived by taking the correlation values in A-B and scaling by their normalised covariance. Note here that the images become more focal, with visual areas highlighted for low phase change and medial frontal cortices highlighted when the phase change approaches 90°. The increased focality of the highlighted regions likely results from the fact that variance normalisation, inherent in the correlation images, amplifies leakage close to the centre of the head; the fact that the variance of such regions is down-weighted by the beamformer means that they fail to appear when covariance is employed.

**Figure 2.**
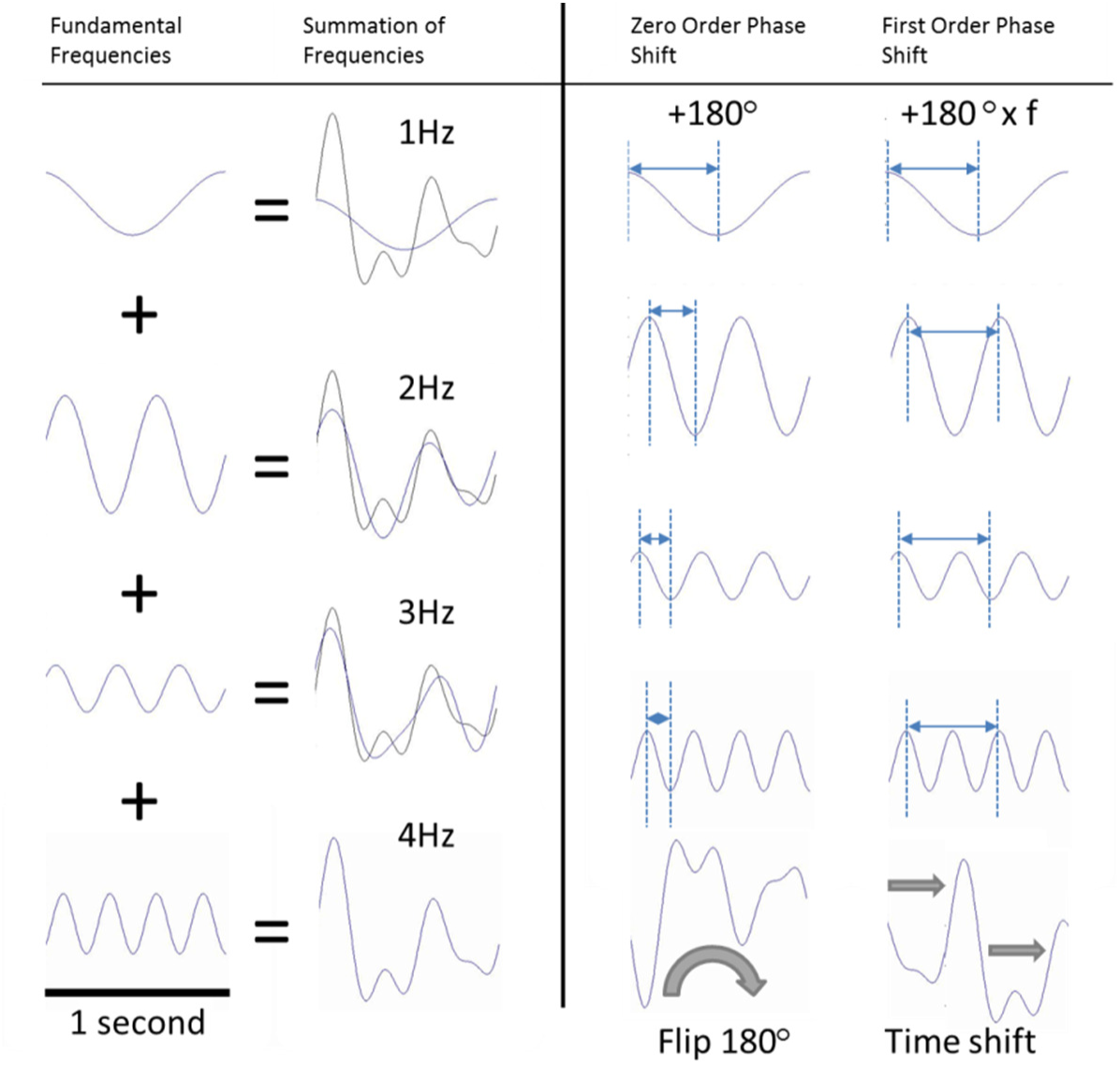
Left panel illustrates how the summation of sine waves at different frequencies and phases results in a signal with a complicated waveform. The right panel illustrates the effect of zero order and first order phase shifting on the same constituent sine waves that make up the complex time series. The ZOPS method results in a rotation of the waveform about the complex axis. If all sine waves are shifted by 180 degrees, this results in a complete inversion of the timecourse. In the FOPS each frequency is shifted by a constant time regardless of frequency, resulting in a simple translation of the wave in the time domain.

Figures 2E and 2F contain example timecourse fits, for voxels located in the frontal cortex. The seed timecourse, *Y*_1_, from the primary visual cortex, is shown in green, the test voxel timecourse, *Y*_2_ is shown in blue and the model fit, 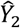, is shown in red. Note the excellent agreement between the model and the data, particularly for the relevant condition. Figure 2G shows phase maps, highlighting the best fitting phase angle for each voxel in the head. The relevant condition is shown on the left and irrelevant on the right. Note the higher phase values in the frontal regions for the relevant condition. Note also the difference between the relevant and irrelevant conditions. Overall, the results presented in Figure 2 show good evidence that the ZOPS model generates good fits to measured theta-band envelope waveforms. Furthermore, the fitted phase angles change with location in the brain, with increased phase occurring in the frontal regions. Note also that the fitted parameters change with task condition.

### Comparison of ZOPS and FOPS

Figure 4 shows a comparison of the zero-order and first-order phase shift methods, for both the relevant and irrelevant condition. Correlation values (derived from the optimum phase angle) are plotted against Euclidean distance between the seed and the test voxel. The ZOPS model is shown in blue, and the FOPS model in red. Figure 4A shows a single point for every voxel whilst in Figure 4B, voxels are grouped into 9 equally spaced bins (representing Euclidean distance) and the mean and standard deviations across voxels in each bin displayed. The primary result shows that the ZOPS method is better able to explain variance in the data compared to the FOPS method for large Euclidean distances in the brain, specifically for those regions greater than ∼ 120mm from the seed. As shown by the inset image, these voxels were primarily located in the frontal lobes. The difference between the two models, for the relevant condition and for voxels >120mm from the seed, was analysed statistically, revealing a significant difference in between methods (*p*<0.05). There was no significant difference between models for the irrelevant condition. The dotted line represents the 95^th^ percentile of the null distribution computed via the monte-carlo phase randomisation procedure (outlined in methods).

### Analysis in Schizophrenia and Correlations with Behaviour

Finally, we used the ZOPS model to look for interrelationships between the visual cortex and the frontal lobes in patients with schizophrenia. Figures 3A-D shows correlation and covariance maps, for the relevant and irrelevant conditions, in a group of 10 patients. Visual comparison between the control group in Figure 1 and the patient group in Figure 3 reveals differences between patients and controls in the magnitude of the values in the frontal cortex. Figures 3E and F show a comparison of the mean measured phase angle, plotted as a function of Euclidean distance from the seed, for the relevant (green) and irrelevant (red) conditions. Note that for group statistical comparisons, ZOPS analysis was performed on individual subjects giving a phase estimate per voxel, per subject. The results, illustrated in figure 3, show that in the control group, task condition had a large modulatory effect on delay. However, in the patient group delay did not appear to be modulated by task condition. For statistical comparison, the last 2 bins (those that encompass the frontal cortices) were entered into a repeated measures ANOVA [2 distance × 2 condition] with group as a between subjects factor, and the mean phase in each bin as the dependent variable. The analysis revealed a significant effect of condition (*F*=7.35, *p*<0.05), a significant effect of distance (*F*=5.09, p<0.05), and a significant interaction between group and condition (*F*=4.87, *p*<0.05). This demonstrates that overall, phase significantly changes with distance from seed location, and the presence of a task induces a significant change in phase in the controls but not the patients.

**Figure 3.**
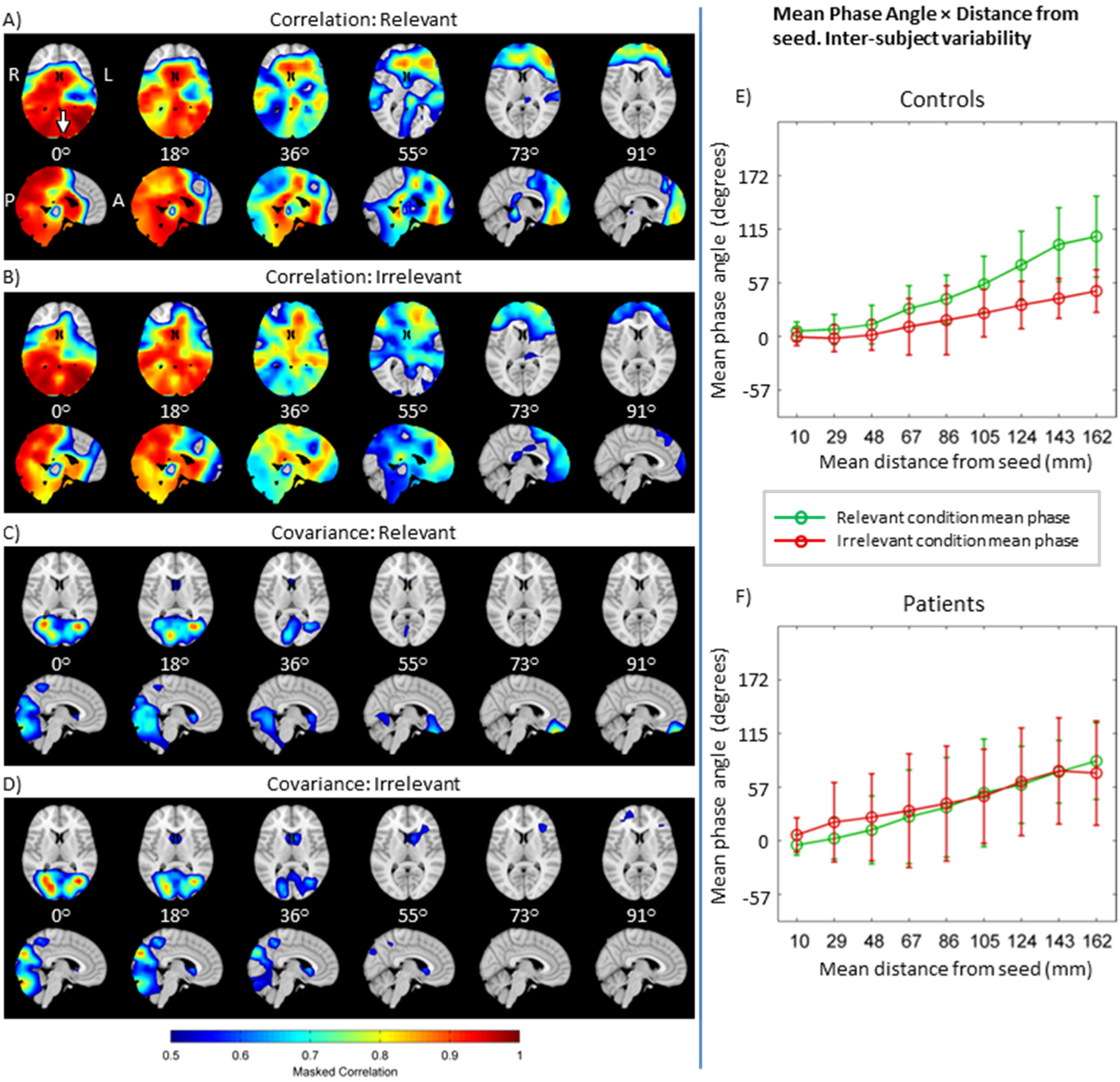
Correlation and covariance masked correlation for schizophrenia patients. Note that the thresholds on these maps have been altered to reveal peak locations in the patient group that were below the 0.5 threshold used in Figure 1. A and B show covariance for the relevant and irrelevant conditions respectively. Likewise, C and D show covariance masked correlation for the relevant and irrelevant conditions respectively. Here the differences between the patients, and controls shown in Figure 1. E-F) Phase change, plotted as a function of Euclidean distance from the seed location, for relevant (green) and irrelevant conditions (red). E) shows controls and F) shows patients – note that the difference induced by task condition in controls is absent in patients.

**Figure 3.**
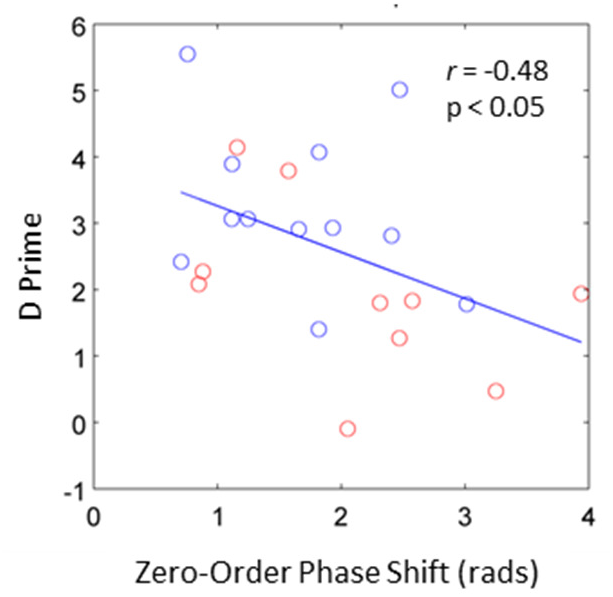
Scatter plot showing the correlation between mean phase estimate in the last distance bin (>140mm) and task performance (D’). To maximise variance, both patients and controls were included. The Y-axis shows D-Prime, a measure of the subject’s ability to detect a target. Phase delay was negatively correlated with D-Prime (r=-0.48, p<0.05) supporting the hypothesis that task performance is related to inter-regional phase delay in the theta BLE.

**Figure 3.**
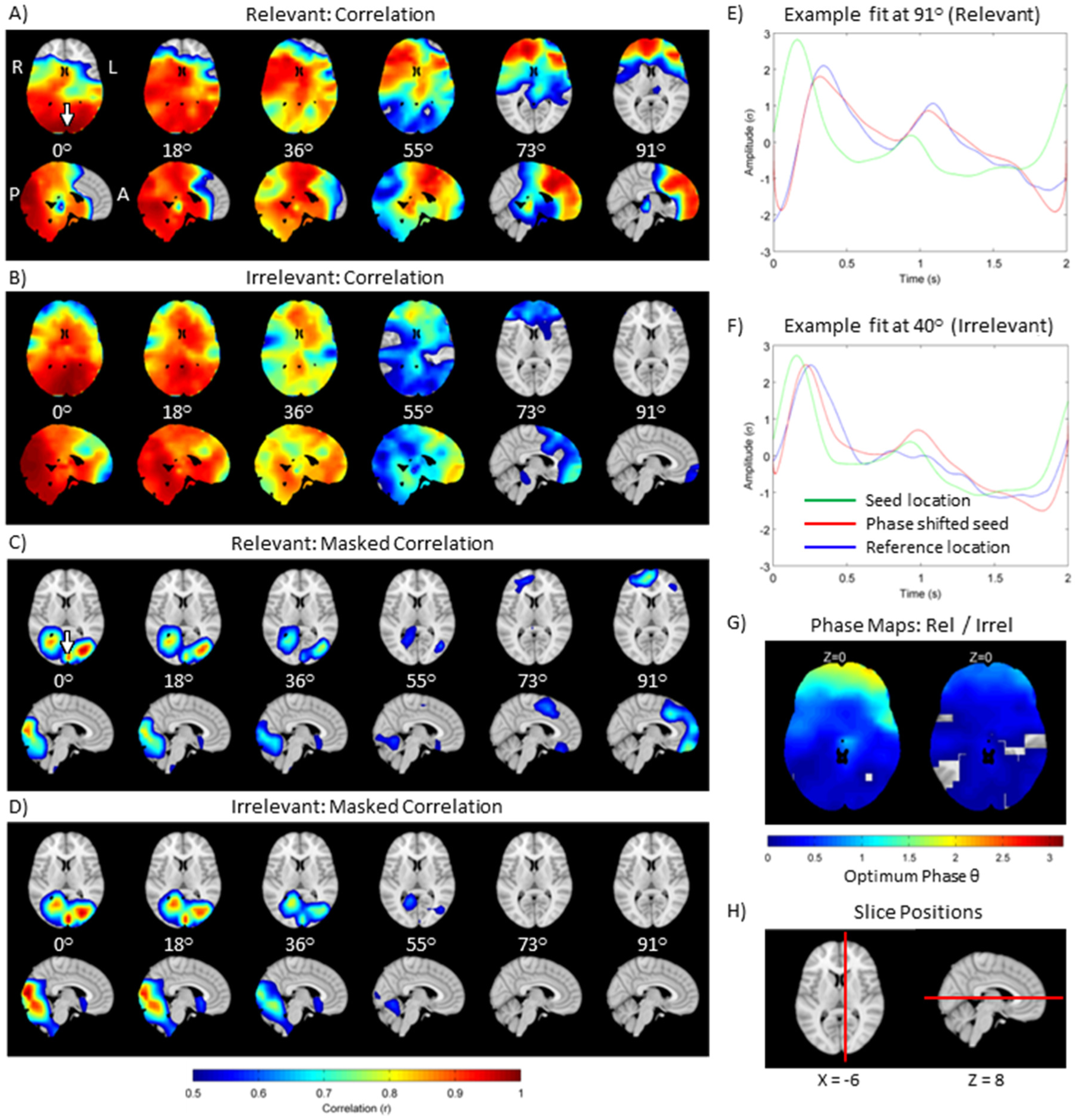
A-B) Correlation images overlaid onto axial and sagittal slices of the standard brain, for 6 linearly spaced phase shifts between 0 and 91 degrees. The lower threshold is set to r = 0.5 for visualisation. A and B show the relevant and irrelevant conditions respectively. C-D) Masked correlation maps, derived by taking the correlation values in A-B and scaling by their normalised covariance. C and D show relevant and irrelevant conditions respectively. Note that scaling by covariance offers more focal peaks in correlation. E-F) Example fits with the seed timecourse, *Y*_1_ shown in green, the test voxel timecourse, *Y*_2_ shown in blue and the model fit, 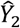, shown in red (E = relevant condition, F = irrelevant condition). G) Phase maps, showing the best fitting phase, in radians, for each voxel. The relevant condition is shown on the left and irrelevant on the right. Note the higher phase values in the frontal regions for the relevant condition. H) Anatomical image showing slice positioning.

**Figure 3).**
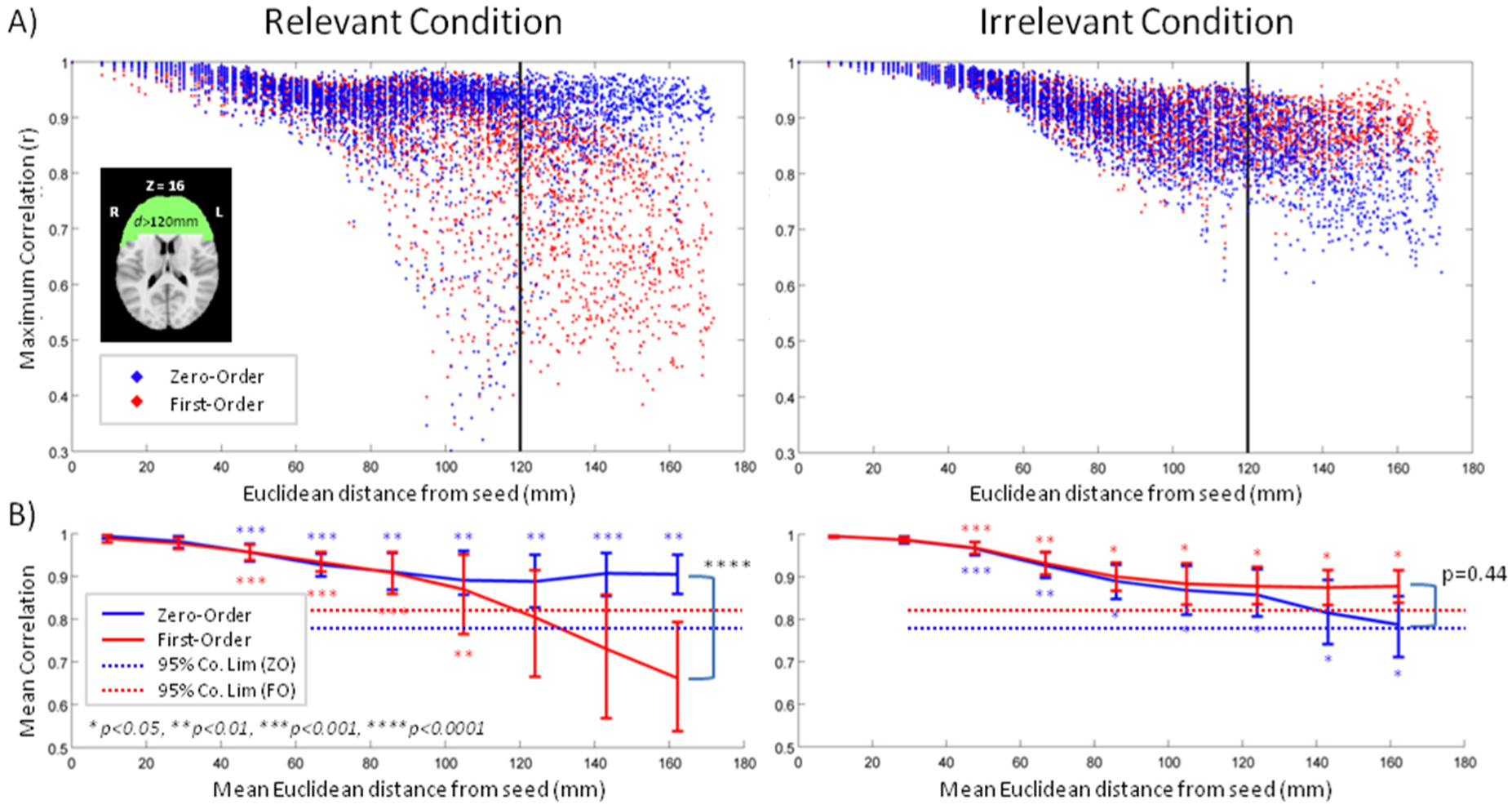
Comparison between the ZO and FO phase shift methods. A) Maximum correlation achieved for each voxel against distance from the seed: ZOPS shown in blue, FOPS in red. B) Mean and standard deviation across voxels for each distance bin. The 95% confidence interval generated for each method is plotted as a dotted line. In both A and B, the relevant condition is shown on the left hand side and irrelevant on the right hand side.

Finally, Figure 4 shows correlation between the measured phase from the ZOPS model, and behaviour; specifically we employed D-Prime as a measure of signal detection on the RM task (see methods). The results show that the degree of phase shift in the frontal cortex, during the relevant condition of the task, negatively correlates with discriminability (*r*=-0.48, *p*<0.05, two tailed).

## Supplementary Figures

**Supp. Fig. 1.**
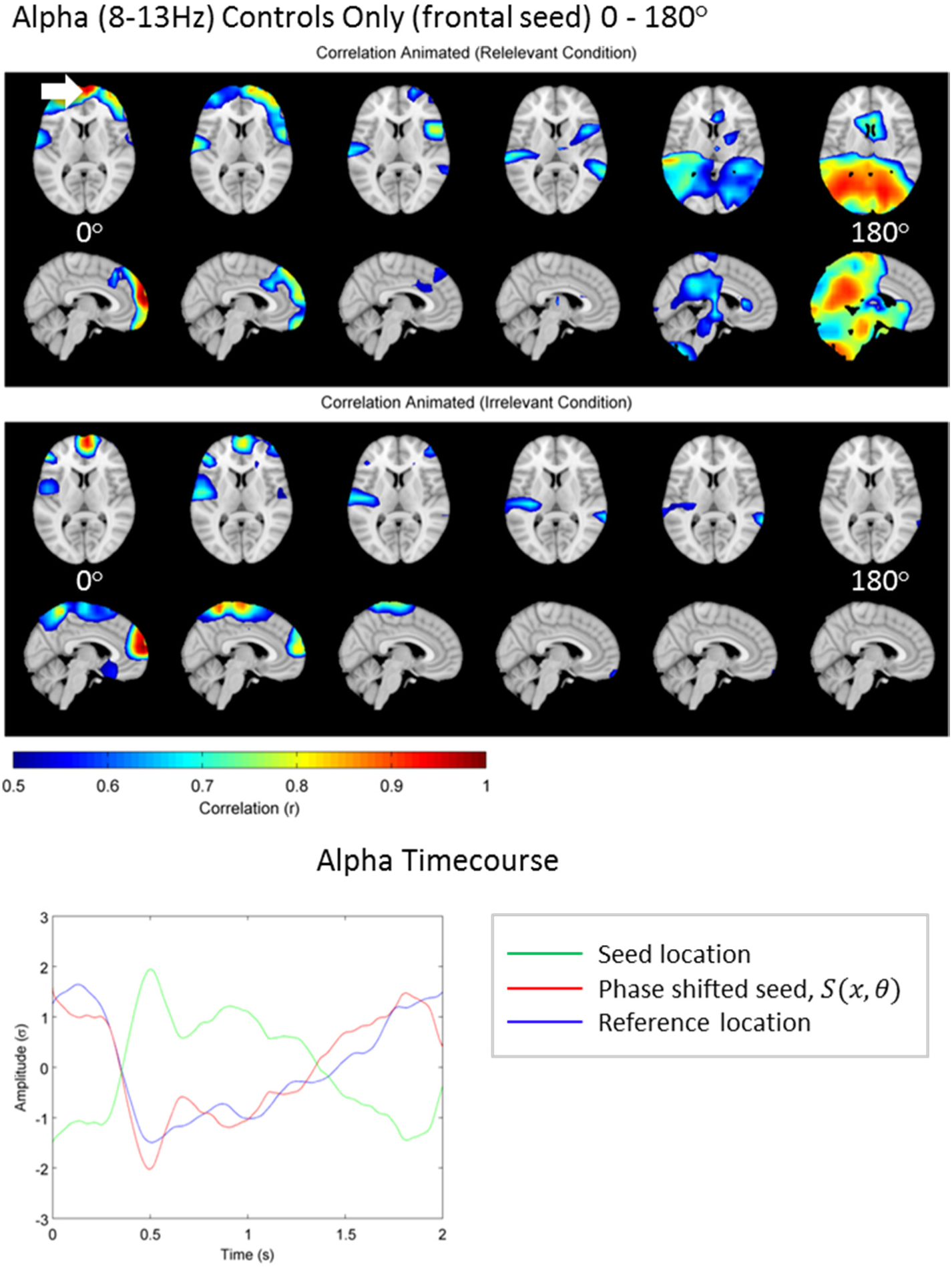
The alpha band appears to show a 180 degree phase shift. i.e. an anti-correlation.

**Supp Fig. 2.**
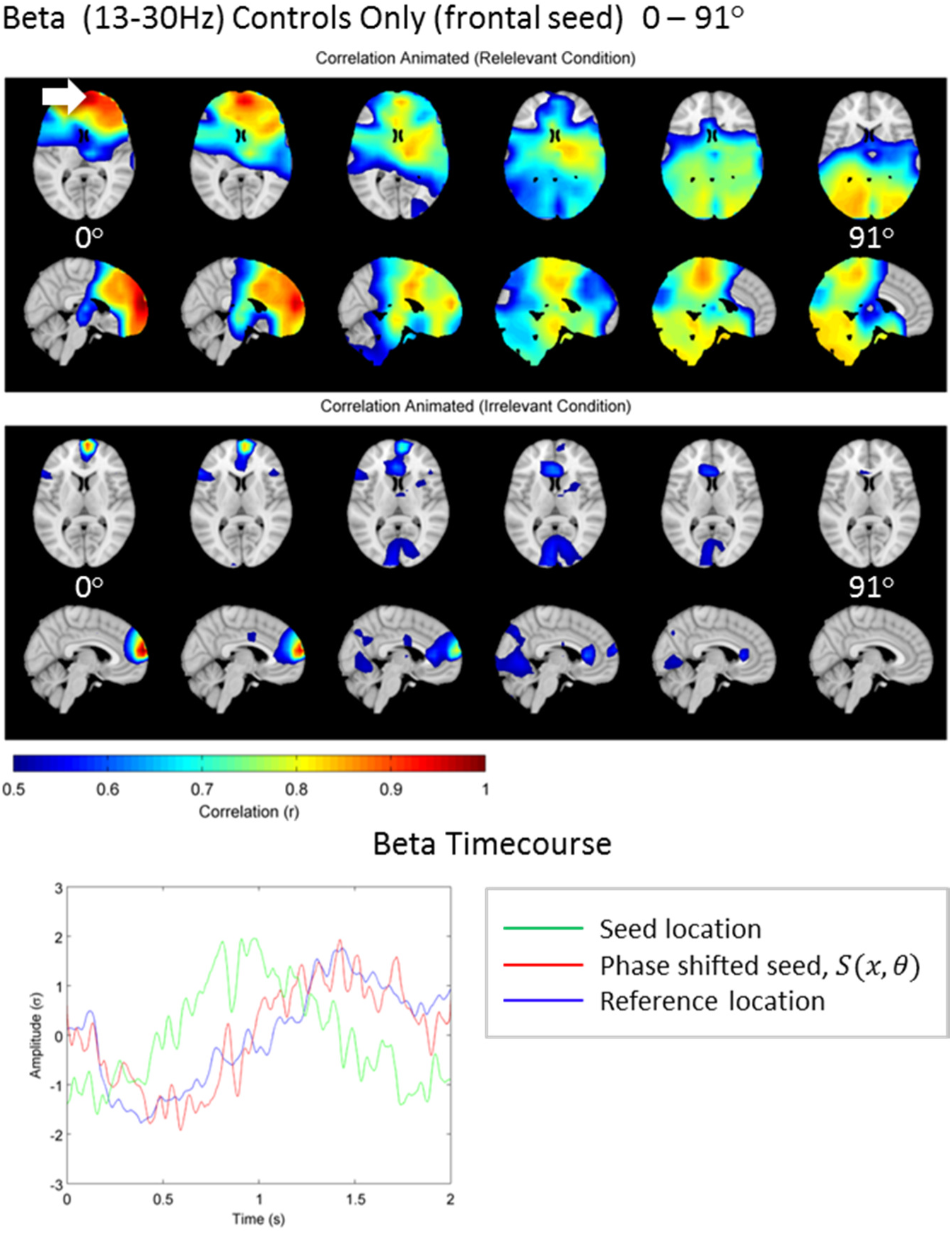
Beta band appears to show, a 90 degree phase lag between frontal and visual cortex. Here the beta-band power fluctuations in frontal cortex appear to precede those in visual cortex.

## Discussion

We have introduced a novel method to assess transient task induced connectivity between brain regions, based upon measurement of a phase delay between theta-band envelope signals measured across the entire cortex. We have compared two separate phase based models; in our first model, termed first order phase shift (FOPS), we assume that all frequency components in the theta envelope will travel through the brain at the same velocity, leading to a time shifted version of the seed envelope at the second brain region. In our second model, termed zero order phase shift (ZOPS), we propose that faster and more efficient processes will manifest in the envelope signal as higher frequency components. These components of the signal will travel with a higher effective velocity than the slower components, leading to a temporally distorted version of the seed envelope at distant brain regions.

We have shown that, at least for the cognitive task employed here, the ZOPS significantly better than the FOPS model, and that the ZOPS model performs significantly better than chance when using randomised surrogate data. Furthermore, we have shown that the fitted phase values are altered significantly by task condition, they are correlated with a behavioural measure, and the condition dependent modulation of phase seen in healthy subjects is not present in patients with schizophrenia. This methodology represents a fundamentally new way to measure directed functional connectivity in the brain.

### Correlation vs. Covariance

Correlation metrics of connectivity used here appeared to reveal that a large proportion of the brain is connected during the relevant condition of the task, with a spatio-temporal pattern similar to a travelling wave of theta-band activity. However, using covariance data in the theta-band allowed the discrimination of a task relevant network involving primarily the occipital lobes and medial frontal cortex. These findings appear to be consistent with previous research implicating theta oscillations in long range connectivity (Buzsaki and Draguhn, 2004; Engel et al., 2001; von Stein and Sarnthein, 2000; Varela et al., 2001) and research relating theta amplitude modulation to task performance in the frontal midline (Brookes et al., 2011b; Gevins et al., 1997; Jensen and Tesche, 2002; Luft et al., 2013). In the present study, mid frontal theta envelope variance is almost entirely explained (R^2^ = 0.94 at the peak) by a ZOPS of the envelope from V1 indicating a close coupling of these two regions. We have also shown that inter-cortical latency is dependent on task condition, which had a significant modulatory effect on the dynamics of the theta envelope. The context dependent modulation of the theta band suggests some task dependent top-down modulation of feed-forward influences, in line with current predictive coding models of brain function and cognition (Arnal and Giraud, 2012; Engel et al., 2001; Friston, 2005). Abnormalities in the establishment of long-range connectivity via low frequency oscillations appear to be disrupted in patients

### Neural Mechanisms Underpinning the ZOPS Model

We have used a mapping function with an occipital seed as the input to accurately model theta-band envelope time-courses across many distant cortical areas. However, it is unclear how the carrier signal of these envelopes supports this close relationship. It is known that connectivity in the envelope can exist without measurable phase coherence in the carrier signal, particularly in long range connections (Bruns et al., 2000). Furthermore, the effective connectivity observed here in the theta band was observed in the group average per condition. Phase coherence must necessarily be measured in the raw signal, to avoid cancellation caused by phase jitter. In future, we plan to use phase-based measures, such as granger causality to investigate the relationship between the carrier signal and the envelope.

### Travelling Waves as Possible Mechanisms of Oscillatory Influence

Although the model put forward here has performed well in describing inter-regional variation in the band-limited envelope, the precise neurobiological mechanisms underpinning the apparent travelling wave observed are not fully understood. Inter-cortical phase coupling is widely thought to be the primary mechanism for communication between cortical sites (Engel et al., 2001; Fries, 2005; Salinas and Sejnowski, 2001) and is modulated by stimulus context, task, or cognitive setting (Engel et al., 2001; Siegel et al., 2012; Singer, 1999). However, our model is based on coupling of signal envelopes rather than oscillatory phase, and as discussed, the two are not directly related. Engel et al. (2013) points out that short periods of non-stationary network connectivity (termed intrinsic coupling modes; or ICMs) may occur between cortical regions either in the oscillatory signal or in the band-limited envelope, and they are both potentially present in the task positive state. Further work is clearly needed to understand the mechanisms responsible for the phenomena observed here, and that should include measures of phase coherence between regions found to have high effective connectivity in the post-stimulus envelope. However, it is possible that measuring the post-stimulus envelope has allowed the measurement of some aspect of brain function that is not normally visible in the oscillatory phase – perhaps due to SNR issues in the oscillatory signal, or because those cortical processes that underlie these envelope modulations do not necessarily rely on phase coupling. We observed clear travelling wave like phenomena in the envelope that shares some similarities with TW phenomena in the oscillatory signal of the EEG and MEG.

It has been shown that cortical travelling exist in the oscillatory signal at varying spatial scales ranging from the columnar scale covering small patches of neurons a few millimetres across (Brookes et al., 2011a; Sato et al., 2012), intra-regional scales, such as within the entire primary visual cortex or motor cortex (Freeman and Barrie, 2000; Rubino et al., 2006; Takahashi et al., 2011) and inter-regional scale covering the whole brain (Alexander et al., 2006; Brookes et al., 2011a, 2011a; Nunez, 1974a; Nunez et al., 1997). Nunez (1974a) introduced the concept of the global theory of standing and travelling waves and proposed the *brain wave equation* as a theoretical model for the global wave like phenomena observed in the EEG. The general postulate put forward (Nunez and Srinivasan, 2006) is that oscillatory waves observed in the EEG are partly composed of TWs, as the term is used in the physical sciences to describe a disturbance that moves through a medium (for example, as sound waves propagate through air). A number of experimental studies have demonstrated the existence of wave like behaviour in the EEG during wakefulness in the alpha band (Nunez, 1974b), slow wave sleep (Massimini et al., 2004, 2007, 2009), and in steady state visually evoked potentials (SSVEPs) (Burkitt et al., 2000; Hughes, 1995; Klimesch et al., 2007; Thorpe et al., 2007), demonstrating the general feasibility of viewing the cortex as a medium through which waves of excitatory and inhibitory action may travel. One major drawback in the experimental study of TWs is that they are degraded by trial averaging of the post stimulus response. In a study employing EEG, ECoG, and MEG, Alexander et al. (2011a) found that travelling waves analysed on a single trial basis accounted for the spatiotemporal pattern of observed ERs in trial-averaged data, but as evoked responses depend mainly on phase locking, trial averaging obscures travelling wave like phenomena. Evoked responses can therefore be explained as TWs and standing waves in the global field oscillations that are phase locked to the stimulus only at specific locations across the cortex, leading to an interference pattern in the trial averaged signal.

Consequently, while evoked responses may be a useful indicator of primary sensory processing latency (Henson and Rugg, 2003), it is likely that trial averaging obscures important information relating to cortical dynamics and stimulus driven connectivity. Those methods described above attempt to overcome this issue by measuring phase differences between adjacent sensors, building up a picture of the phase gradient (2011a, 2011a). The phase gradient therefore shows the general direction of wave propagation by indexing the phase coherence between adjacent sensors. However, phase gradients do not necessarily lead to long-range coherence as cumulative phase jitter may result in phase randomisation at long distances. Alexander found that the theta band TW moves in the posterior-anterior direction while alpha moves in the anterior-posterior direction, which is broadly in agreement with both Bastos (Bastos et al., 2015a) and our results.

Research into travelling waves outlined above points to the possibility that the evoked response and the induced response are closely related; the induced response is simply a time shifted version of the evoked response with randomised phase in the carrier signal. In fact, there is some evidence to suggest that components of the evoked response that are lost in trial averaging of the raw signal due to phase variations may be transferred to the induced response (David et al., 2005; Makeig, 1993). These two aspects of post stimulus activity (evoked and induced) are therefore closely related and may together form a travelling wave like pattern in the envelope signal (that contains both evoked and induced responses) as the later cortical events occur at more distal regions from the origin. In our case, the evoked response in the visual cortex marks the origin of a travelling wave that becomes increasingly phase incoherent as it moves further away from the origin. It is therefore concluded that temporal delay observed between cortical sites involves the gradual transition from a phase locked to a nonphase locked response, which is only visible in the envelope signal. This idea is directly testable by measuring how the evoked signal relates to the travelling wave in the envelope observed here and could be the focus of further investigations.

## References

Abeles, M. (1982). Role of the cortical neuron: integrator or coincidence detector? Isr. J. Med. Sci. 18, 83–92.

Alexander, D.M., Arns, M.W., Paul, R.H., Rowe, D.L., Cooper, N., Esser, A.H., Fallahpour, K., Stephan, B.C., Heesen, E., and Breteler, R. (2006). EEG markers for cognitive decline in elderly subjects with subjective memory complaints. J. Integr. Neurosci. 5, 49–74.

Alonso, J.-M., Usrey, W.M., and Reid, R.C. (1996). Precisely correlated firing in cells of the lateral geniculate nucleus. Nature 383, 815–819.

Ammons, R.B., and Ammons, C.H. (1962). The Quick Test (QT): Provisional manual. Psychol. Rep. 11, 111–161.

Arnal, L.H., and Giraud, A.-L. (2012). Cortical oscillations and sensory predictions. Trends Cogn. Sci. 16, 390–398.

Azouz, R., and Gray, C.M. (2000). Dynamic spike threshold reveals a mechanism for synaptic coincidence detection in cortical neurons in vivo. Proc. Natl. Acad. Sci. 97, 8110–8115.

Bastos, A.M., Vezoli, J., Bosman, C.A., Schoffelen, J.-M., Oostenveld, R., Dowdall, J.R., De Weerd, P., Kennedy, H., and Fries, P. (2015a). Visual areas exert feedforward and feedback influences through distinct frequency channels. Neuron 85, 390–401.

Bastos, A.M., Litvak, V., Moran, R., Bosman, C.A., Fries, P., and Friston, K.J. (2015b). A DCM study of spectral asymmetries in feedforward and feedback connections between visual areas V1 and V4 in the monkey. Neuroimage 108, 460–475.

Brookes, M.J., Hale, J.R., Zumer, J.M., Stevenson, C.M., Francis, S.T., Barnes, G.R., Owen, J.P., Morris, P.G., and Nagarajan, S.S. (2011a). Measuring functional connectivity using MEG: methodology and comparison with fcMRI. Neuroimage 56, 1082–1104.

Brookes, M.J., Wood, J.R., Stevenson, C.M., Zumer, J.M., White, T.P., Liddle, P.F., and Morris, P.G. (2011b). Changes in brain network activity during working memory tasks: A magnetoencephalography study. NeuroImage 55, 1804–1815.

Brookes, M.J., Woolrich, M.W., and Barnes, G.R. (2012). Measuring functional connectivity in MEG: a multivariate approach insensitive to linear source leakage. Neuroimage 63, 910–920.

Bruno, R.M., and Sakmann, B. (2006). Cortex is driven by weak but synchronously active thalamocortical synapses. Science 312, 1622–1627.

Bruns, A., Eckhorn, R., Jokeit, H., and Ebner, A. (2000). Amplitude envelope correlation detects coupling among incoherent brain signals. Neuroreport 11, 1509–1514.

Burkitt, G.R., Silberstein, R.B., Cadusch, P.J., and Wood, A.W. (2000). Steady-state visual evoked potentials and travelling waves. Clin. Neurophysiol. 111, 246–258.

Buzsaki, G., and Draguhn, A. (2004). Neuronal oscillations in cortical networks. Science 304, 1926–1929.

David, O., Harrison, L., and Friston, K.J. (2005). Modelling event-related responses in the brain. NeuroImage 25, 756–770.

Donner, T.H., Sagi, D., Bonneh, Y.S., and Heeger, D.J. (2008). Opposite neural signatures of motion-induced blindness in human dorsal and ventral visual cortex. J. Neurosci. 28, 10298–10310.

Donner, T.H., Siegel, M., Fries, P., and Engel, A.K. (2009). Buildup of choice-predictive activity in human motor cortex during perceptual decision making. Curr. Biol. 19, 1581–1585.

Driver, J., Blankenburg, F., Bestmann, S., Vanduffel, W., and Ruff, C.C. (2009). Concurrent brain-stimulation and neuroimaging for studies of cognition. Trends Cogn. Sci. 13, 319–327.

Eckhorn, R., Bauer, R., Jordan, W., Brosch, M., Kruse, W., Munk, M., and Reitboeck, H.J. (1988). Coherent oscillations: A mechanism of feature linking in the visual cortex? Biol. Cybern. 60, 121–130.

Engel, A.K., and Singer, W. (2001). Temporal binding and the neural correlates of sensory awareness. Trends Cogn Sci 5, 16–25.

Engel, A.K., König, P., and Singer, W. (1991). Direct physiological evidence for scene segmentation by temporal coding. Proc. Natl. Acad. Sci. 88, 9136–9140.

Engel, A.K., Fries, P., and Singer, W. (2001). Dynamic predictions: Oscillations and synchrony in top–down processing. Nat. Rev. Neurosci. 2, 704–716.

Engel, A.K., Gerloff, C., Hilgetag, C.C., and Nolte, G. (2013). Intrinsic coupling modes: multiscale interactions in ongoing brain activity. Neuron 80, 867–886.

Felleman, D.J., and Van Essen, D.C. (1991). Distributed hierarchical processing in the primate cerebral cortex. Cereb. Cortex 1, 1–47.

Freeman, W.J., and Barrie, J.M. (2000). Analysis of spatial patterns of phase in neocortical gamma EEGs in rabbit. J. Neurophysiol. 84.

Fries, P. (2005). A mechanism for cognitive dynamics: neuronal communication through neuronal coherence. Trends Cogn. Sci. 9, 474–480.

Friston, K. (2005). A theory of cortical responses. Philos. Trans. R. Soc. B Biol. Sci. 360, 815–836.

Friston, K.J. (1999). Schizophrenia and the disconnection hypothesis. Acta Psychiatr. Scand. Suppl. 395, 68–79.

Friston, K.J., Harrison, L., and Penny, W. (2003). Dynamic causal modelling. NeuroImage 19, 1273–1302.

Gevins, A., Smith, M.E., McEvoy, L., and Yu, D. (1997). High-resolution EEG mapping of cortical activation related to working memory: effects of task difficulty, type of processing, and practice. Cereb. Cortex 7, 374–385.

Glimcher, P.W. (2003). The neurobiology of visual-saccadic decision making. Annu. Rev. Neurosci. 26, 133–179.

Gold, J.I., and Shadlen, M.N. (2001). Neural computations that underlie decisions about sensory stimuli. Trends Cogn. Sci. 5, 10–16.

Gow Jr, D.W., Segawa, J.A., Ahlfors, S.P., and Lin, F.-H. (2008). Lexical influences on speech perception: a Granger causality analysis of MEG and EEG source estimates. Neuroimage 43, 614–623.

Gray, C.M., and Singer, W. (1989). Stimulus-specific neuronal oscillations in orientation columns of cat visual cortex. Proc. Natl. Acad. Sci. 86, 1698–1702.

Gray, C.M., König, P., Engel, A.K., and Singer, W. (1989). Oscillatory responses in cat visual cortex exhibit inter-columnar synchronization which reflects global stimulus properties. Nature 338, 334–337.

Gregoriou, G.G., Gotts, S.J., Zhou, H., and Desimone, R. (2009). High-Frequency, Long-Range Coupling Between Prefrontal and Visual Cortex During Attention. Science 324, 1207–1210.

Gross, J., Kujala, J., Hamalainen, M., Timmermann, L., Schnitzler, A., and Salmelin, R. (2001). Dynamic imaging of coherent sources: Studying neural interactions in the human brain. Proc. Natl. Acad. Sci. U. S. A. 98, 694–699.

Hall, E.L., Woolrich, M.W., Thomaz, C.E., Morris, P.G., and Brookes, M.J. (2013). Using variance information in magnetoencephalography measures of functional connectivity. NeuroImage 67, 203–212.

Henson, R.N.A., and Rugg, M.D. (2003). Neural response suppression, haemodynamic repetition effects, and behavioural priming. Neuropsychologia 41, 263–270.

Hipp, J.F., Hawellek, D.J., Corbetta, M., Siegel, M., and Engel, A.K. (2012). Large-scale cortical correlation structure of spontaneous oscillatory activity. Nat. Neurosci. 15, 884–890.

Horwitz, G.D., Batista, A.P., and Newsome, W.T. (2004). Representation of an abstract perceptual decision in macaque superior colliculus. J. Neurophysiol. 91, 2281–2296.

Huang, M.X., Mosher, J.C., and Leahy, R.M. (1999). A sensor-weighted overlapping-sphere head model and exhaustive head model comparison for MEG. Phys. Med. Biol. 44, 423–440.

Hughes, J.R. (1995). The phenomenon of travelling waves: a review. Clin. EEG Neurosci. 26, 1–6.

Hutchison, W.D., Dostrovsky, J.O., Walters, J.R., Courtemanche, R., Boraud, T., Goldberg, J., and Brown, P. (2004). Neuronal Oscillations in the Basal Ganglia and Movement Disorders: Evidence from Whole Animal and Human Recordings. J. Neurosci. 24, 9240–9243.

Ioannides, A.A., Liu, L.C., Kwapien, J., Drozdz, S., and Streit, M. (2000). Coupling of regional activations in a human brain during an object and face affect recognition task. Hum. Brain Mapp. 11, 77–92.

Jenkinson, M., and Smith, S. (2001). A global optimisation method for robust affine registration of brain images. Med. Image Anal. 5, 143–156.

Jensen, O., and Tesche, C.D. (2002). Frontal theta activity in humans increases with memory load in a working memory task. Eur. J. Neurosci. 15, 1395–1399.

Jerbi, K., Lachaux, J.-P., Karim, N., Pantazis, D., Leahy, R.M., Garnero, L., and Baillet, S. (2007). Coherent neural representation of hand speed in humans revealed by MEG imaging. Proc. Natl. Acad. Sci. 104, 7676–7681.

Kastner, S., and Ungerleider, L.G. (2000). Mechanisms of visual attention in the human cortex. Annu. Rev. Neurosci. 23, 315–341.

Kim, J.-N., and Shadlen, M.N. (1999). Neural correlates of a decision in the dorsolateral prefrontal cortex of the macaque. Nat. Neurosci. 2, 176–185.

Klimesch, W., Hanslmayr, S., Sauseng, P., Gruber, W.R., and Doppelmayr, M. (2007). P1 and traveling alpha waves: evidence for evoked oscillations. J. Neurophysiol. 97, 1311–1318.

König, P., Engel, A.K., and Singer, W. (1996). Integrator or coincidence detector? The role of the cortical neuron revisited. Trends Neurosci. 19, 130–137.

Leckman, J.F., Sholomskas, D., Thompson, W.D., Belanger, A., and Weissman, M.M. (1982). Best estimate of lifetime psychiatric diagnosis: a methodological study. Arch. Gen. Psychiatry 39, 879–883.

Leopold, D.A., Murayama, Y., and Logothetis, N.K. (2003). Very slow activity fluctuations in monkey visual cortex: implications for functional brain imaging. Cereb. Cortex 13, 422–433.

Liddle, E.B., Hollis, C., Batty, M.J., Groom, M.J., Totman, J.J., Liotti, M., Scerif, G., and Liddle, P.F. (2011). Task-related default mode network modulation and inhibitory control in ADHD: effects of motivation and methylphenidate. J. Child Psychol. Psychiatry 52, 761–771.

Liddle, E.B., Price, D., Palaniyappan, L., Brookes, M.J., Robson, S.E., Hall, E.L., Morris, P.G., and Liddle, P.F. (2016). Abnormal salience signaling in schizophrenia: The role of integrative beta oscillations. Hum. Brain Mapp.

Liddle, P.F., Ngan, E.T.C., Duffield, G., Kho, K., and Warren, A.J. (2002). Signs and Symptoms of Psychotic Illness (SSPI): a rating scale. Br. J. Psychiatry J. Ment. Sci. 180, 45–50.

Liu, Z., Fukunaga, M., de Zwart, J.A., and Duyn, J.H. (2010). Large-scale spontaneous fluctuations and correlations in brain electrical activity observed with magnetoencephalography. Neuroimage 51, 102–111.

Luft, C.D.B., Nolte, G., and Bhattacharya, J. (2013). High-learners present larger mid-frontal theta power and connectivity in response to incorrect performance feedback. J. Neurosci. 33, 2029–2038.

Makeig, S. (1993). Auditory event-related dynamics of the EEG spectrum and effects of exposure to tones. Electroencephalogr. Clin. Neurophysiol. 86, 283–293.

Markov, N.T., Vezoli, J., Chameau, P., Falchier, A., Quilodran, R., Huissoud, C., Lamy, C., Misery, P., Giroud, P., Ullman, S., et al. (2014). Anatomy of hierarchy: Feedforward and feedback pathways in macaque visual cortex. J. Comp. Neurol. 522, 225–259.

Marzetti, L., Della Penna, S., Snyder, A.Z., Pizzella, V., Nolte, G., de Pasquale, F., Romani, G.L., and Corbetta, M. (2013). Frequency specific interactions of MEG resting state activity within and across brain networks as revealed by the multivariate interaction measure. Neuroimage 79, 172–183.

Massimini, M., Huber, R., Ferrarelli, F., Hill, S., and Tononi, G. (2004). The sleep slow oscillation as a traveling wave. J. Neurosci. 24, 6862–6870.

Massimini, M., Ferrarelli, F., Esser, S.K., Riedner, B.A., Huber, R., Murphy, M., Peterson, M.J., and Tononi, G. (2007). Triggering sleep slow waves by transcranial magnetic stimulation. Proc. Natl. Acad. Sci. 104, 8496–8501.

Massimini, M., Tononi, G., and Huber, R. (2009). Slow waves, synaptic plasticity and information processing: insights from transcranial magnetic stimulation and high-density EEG experiments. Eur. J. Neurosci. 29, 1761–1770.

Mazaheri, A., Nieuwenhuis, I.L., van Dijk, H., and Jensen, O. (2009). Prestimulus alpha and mu activity predicts failure to inhibit motor responses. Hum. Brain Mapp. 30, 1791–1800.

Mazaheri, A., Coffey-Corina, S., Mangun, G.R., Bekker, E.M., Berry, A.S., and Corbett, B.A. (2010). Functional disconnection of frontal cortex and visual cortex in attention-deficit/hyperactivity disorder. Biol. Psychiatry 67, 617–623.

Moore, T., and Armstrong, K.M. (2003). Selective gating of visual signals by microstimulation of frontal cortex. Nature 421, 370–373.

Munk, M.H., Roelfsema, P.R., König, P., Engel, A.K., and Singer, W. (1996). Role of reticular activation in the modulation of intracortical synchronization. Science 272, 271–274.

Neuper, C., Wortz, M., and Pfurtscheller, G. (2006). ERD/ERS patterns reflecting sensorimotor activation and deactivation. Prog Brain Res 159, 211–222.

Nienborg, H., and Cumming, B.G. (2009). Decision-related activity in sensory neurons reflects more than a neuron’s causal effect. Nature 459, 89–92.

Nolte, G., Bai, O., Wheaton, L., Mari, Z., Vorbach, S., and Hallett, M. (2004). Identifying true brain interaction from EEG data using the imaginary part of coherency. Clin. Neurophysiol. 115, 2292–2307.

Nunez, P.L. (1974a). The brain wave equation: a model for the EEG. Math. Biosci. 21, 279–297.

Nunez, P.L. (1974b). Wavelike properties of the alpha rhythm. Biomed. Eng. IEEE Trans. On 473–482.

Nunez, P.L., and Srinivasan, R. (2006). A theoretical basis for standing and traveling brain waves measured with human EEG with implications for an integrated consciousness. Clin. Neurophysiol. 117, 2424–2435.

Nunez, P.L., Srinivasan, R., Westdorp, A.F., Wijesinghe, R.S., Tucker, D.M., Silberstein, R.B., and Cadusch, P.J. (1997). EEG coherency: I: statistics, reference electrode, volume conduction, Laplacians, cortical imaging, and interpretation at multiple scales. Electroencephalogr. Clin. Neurophysiol. 103, 499–515.

de Pasquale, F., Della Penna, S., Snyder, A.Z., Lewis, C., Mantini, D., Marzetti, L., Belardinelli, P., Ciancetta, L., Pizzella, V., Romani, G.L., et al. (2010). Temporal dynamics of spontaneous MEG activity in brain networks. Proc. Natl. Acad. Sci. 107, 6040–6045.

Pevalin, D., and Rose, D. (2002). The national statistics socio-economic classification: unifying official and sociological approaches to the conceptualisation and measurement of social class in the United Kingdom. Sociétés Contemp. n^°^ 45-46, 75–106.

Pfurtscheller, G., and Aranibar, A. (1977). Event-related cortical desynchronization detected by power measurements of scalp EEG. Electroencephalogr. Clin. Neurophysiol. 42, 817–826.

Pfurtscheller, G., and Lopes da Silva, F.H. (1999). Event-related EEG/MEG synchronization and desynchronization: basic principles. Clin. Neurophysiol. 110, 1842–1857.

Phillips, W.A., and Silverstein, S.M. (2003). Convergence of biological and psychological perspectives on cognitive coordination in schizophrenia. Behav. Brain Sci. 26, 65–82.

Prichard, D., and Theiler, J. (1994). Generating surrogate data for time series with several simultaneously measured variables. Phys Rev Lett 73, 951–954.

Ress, D., and Heeger, D.J. (2003). Neuronal correlates of perception in early visual cortex. Nat. Neurosci. 6, 414–420.

Robinson, S., and Vrba, J. (1999). Functional neuroimaging by synthetic aperture magnetometry (SAM). In Recent Advances in Biomagnetism, T. Yoshimoto, M. Kotani, S. Kuriki, H. Karibe, and N. Nakasato, eds. (Sendai: Tohoku University Press, Japan), pp. 302–305.

Rubino, D., Robbins, K.A., and Hatsopoulos, N.G. (2006). Propagating waves mediate information transfer in the motor cortex. Nat. Neurosci. 9, 1549–1557.

Salinas, E., and Sejnowski, T.J. (2001). Correlated neuronal activity and the flow of neural information. Nat. Rev. Neurosci. 2, 539–550.

Sarnthein, J., Petsche, H., Rappelsberger, P., Shaw, G.L., and Von Stein, A. (1998). Synchronization between prefrontal and posterior association cortex during human working memory. Proc. Natl. Acad. Sci. 95, 7092–7096.

Sarvas, J. (1987). Basic mathematical and electromagnetic concepts of the biomagnetic inverse problem. Phys. Med. Biol. 32, 11–22.

Sato, T.K., Nauhaus, I., and Carandini, M. (2012). Traveling waves in visual cortex. Neuron 75, 218–229.

Schlögl, A., and Supp, G. (2006). Analyzing event-related EEG data with multivariate autoregressive parameters. Prog. Brain Res. 159, 135–147.

Schnitzler, A., and Gross, J. (2005). Normal and pathological oscillatory communication in the brain. Nat. Rev. Neurosci. 6, 285–296.

Schoffelen, J.-M., and Gross, J. (2009). Source connectivity analysis with MEG and EEG. Hum. Brain Mapp. 30, 1857–1865.

Schölvinck, M.L., Maier, A., Ye, F.Q., Duyn, J.H., and Leopold, D.A. (2010). Neural basis of global resting-state fMRI activity. Proc. Natl. Acad. Sci. 107, 10238–10243.

Serences, J.T., and Yantis, S. (2006). Selective visual attention and perceptual coherence. Trends Cogn. Sci. 10, 38–45.

Siegel, M., Donner, T.H., and Engel, A.K. (2012). Spectral fingerprints of large-scale neuronal interactions. Nat. Rev. Neurosci. 13, 121–134.

Singer, W. (1999). Neuronal synchrony: a versatile code for the definition of relations? Neuron 24, 49–65.

Singer, W., and Gray, C.M. (1995). Visual feature integration and the temporal correlation hypothesis. Annu. Rev. Neurosci. 18, 555–586.

Smith, S.M. (2002). Fast robust automated brain extraction. Hum. Brain Mapp. 17, 143–155.

von Stein, A., and Sarnthein, J. (2000). Different frequencies for different scales of cortical integration: from local gamma to long range alpha/theta synchronization. Int. J. Psychophysiol. 38, 301–313.

von Stein, A., Chiang, C., and König, P. (2000). Top-down processing mediated by interareal synchronization. Proc. Natl. Acad. Sci. 97, 14748–14753.

Stephan, K.E., Harrison, L.M., Kiebel, S.J., David, O., Penny, W.D., and Friston, K.J. (2007). Dynamic causal models of neural system dynamics: current state and future extensions. J. Biosci. 32, 129–144.

Stephane, M., Ince, N.F., Leuthold, A., Pellizzer, G., Tewfik, A.H., Surerus, C., Kuskowski, M., and McClannahan, K. (2008). Temporospatial characterization of brain oscillations (TSCBO) associated with subprocesses of verbal working memory in schizophrenia. Clin EEG Neurosci 39, 194–202.

Takahashi, K., Saleh, M., Penn, R.D., and Hatsopoulos, N.G. (2011). Propagating waves in human motor cortex. Front. Hum. Neurosci. 5.

Tass, P., Rosenblum, M.G., Weule, J., Kurths, J., Pikovsky, A., Volkmann, J., Schnitzler, A., and Freund, H.-J. (1998). Detection of n: m phase locking from noisy data: application to magnetoencephalography. Phys. Rev. Lett. 81, 3291.

Thorpe, S.G., Nunez, P.L., and Srinivasan, R. (2007). Identification of wave-like spatial structure in the SSVEP: Comparison of simultaneous EEG and MEG. Stat. Med. 26, 3911–3926.

Timmermann, L., Wojtecki, L., Gross, J., Lehrke, R., Voges, J., Maarouf, M., Treuer, H., Sturm, V., and Schnitzler, A. (2004). Ten-Hertz stimulation of subthalamic nucleus deteriorates motor symptoms in Parkinson’s disease. Mov. Disord. 19, 1328–1333.

Uhlhaas, P.J., and Singer, W. (2010). Abnormal neural oscillations and synchrony in schizophrenia. Nat Rev Neurosci 11, 100–113.

Uhlhaas, P.J., Roux, F., Rodriguez, E., Rotarska-Jagiela, A., and Singer, W. (2010). Neural synchrony and the development of cortical networks. Trends Cogn. Sci. 14, 72–80.

Van Veen, B.D., Van Drongelen, W., Yuchtman, M., and Suzuki, A. (1997). Localization of brain electrical activity via linearly constrained minimum variance spatial filtering. IEEE Trans. Biomed. Eng. 44, 867–880.

Varela, F., Lachaux, J.-P., Rodriguez, E., and Martinerie, J. (2001). The brainweb: phase synchronization and large-scale integration. Nat. Rev. Neurosci. 2, 229–239.

Womelsdorf, T., Schoffelen, J.-M., Oostenveld, R., Singer, W., Desimone, R., Engel, A.K., and Fries, P. (2007). Modulation of neuronal interactions through neuronal synchronization. Science 316, 1609–1612.

Zanto, T.P., Rubens, M.T., Thangavel, A., and Gazzaley, A. (2011). Causal role of the prefrontal cortex in top-down modulation of visual processing and working memory. Nat. Neurosci. 14, 656–661.

